# Mechanosignaling promotes macrophage apoptosis resistance in pulmonary fibrosis *via* metabolic reprogramming

**DOI:** 10.64898/2026.08.23.746574

**Authors:** Chao He, Cristian Coarfa, Neftali Garcia, Olubunmi C. Lebimoyo, Haiwei Gu, Elisa Ruiz-Echartea, Xiaoli Ji, Alan Waich, Juan D. Zuluaga, Lindsay J. Celada, Scott A. Ochsner, Neil J. McKenna, Jennifer L. Larson-Casey, Sandeep K. Agarwal, Farrah Kheradmand, Yong Zhou, A. Brent Carter, Ivan O. Rosas

## Abstract

The mechanisms underlying disease progression in idiopathic pulmonary fibrosis (IPF) and other interstitial lung diseases remain unclear. Increased extracellular matrix stiffness is a hallmark of fibrotic lung diseases, while monocyte-derived macrophages promote fibrosis progression. However, there is limited understanding if mechanical properties of the fibrotic microenvironment influence macrophage phenotypes and fibrogenesis. Profibrotic macrophages are resistant to apoptosis, which is modulated by enhanced mitochondrial bioenergetics. The objective of this study is to determine how lung tissue stiffness impacts macrophage phenotypes and fibrotic progression. We demonstrate that mechanoactivated macrophages exhibit apoptosis-resistance, increased expression of the antiapoptotic protein Bcl-xL, and elevated mitochondrial oxidative phosphorylation. Critically, the metabolic reprogramming observed in mechanoactivated macrophages is dependent on increased glutaminolysis. Inhibition of glutaminolysis attenuated apoptosis resistance in mechanoactivated macrophages. Moreover, inhibition of Bcl-xL *in vivo* protected mice against experimental pulmonary fibrosis. Lastly, mechanoactivated macrophages produce more profibrotic cytokines and promote extracellular matrix production in precision-cut lung slices. We describe a mechanism by which extracellular matrix stiffness mediates macrophage apoptosis resistance and metabolic reprogramming. Our results identify mechanoactivated apoptosis-resistant macrophages as pro-fibrotic mediators, suggesting a novel therapeutic target in IPF and related fibrotic disorders.

**GRAPHIC ABSTRACT:** 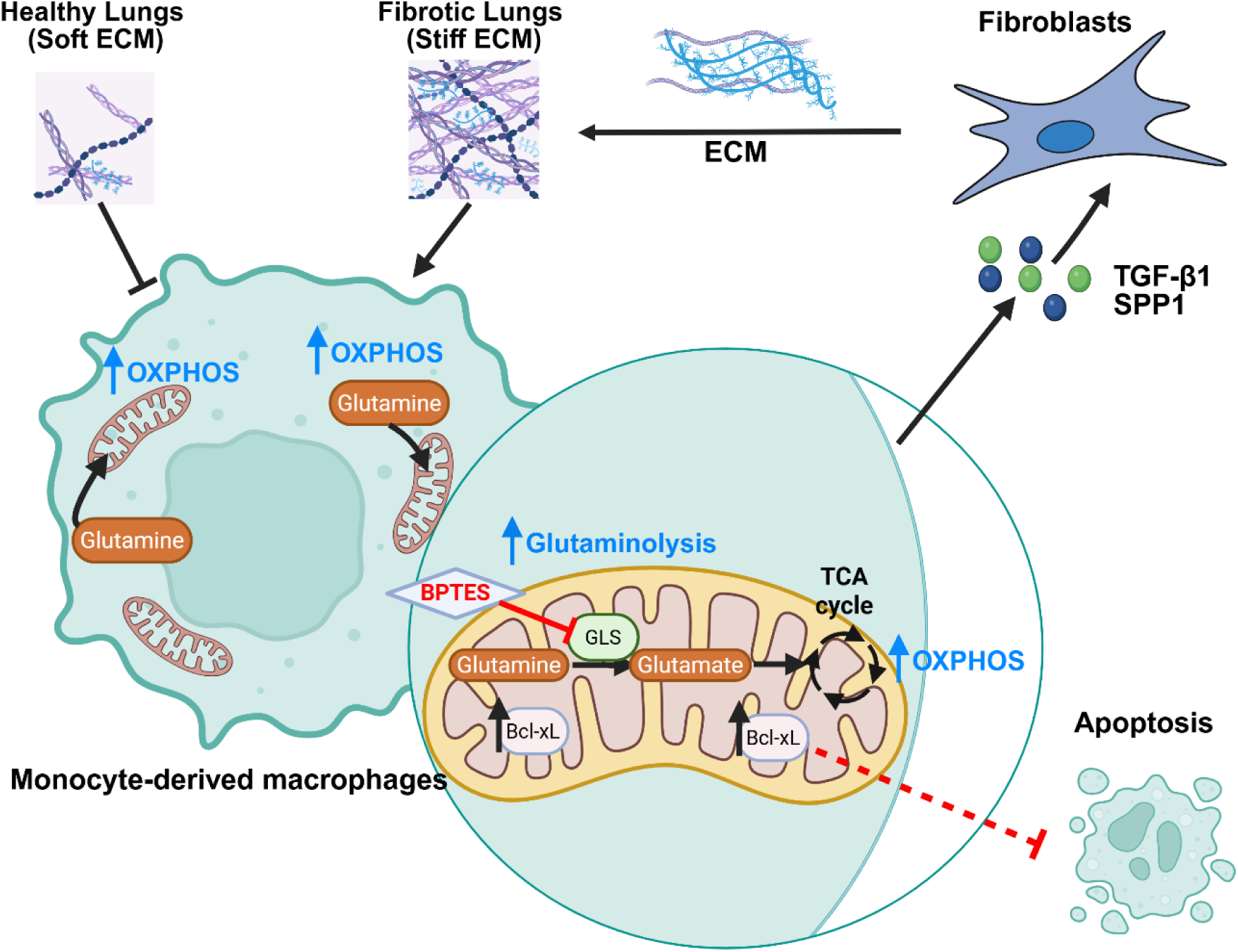

## INTRODUCTION

Idiopathic pulmonary fibrosis (IPF) is a progressive and ultimately fatal interstitial lung disease characterized by chronic scarring of the lung parenchyma, irreversible remodeling of the distal lung, and respiratory failure. IPF has a prevalence approaching 1 in 150 adults aged 65 years or older in the United States, with an even higher prevalence in specific populations such as veterans (1–4). The median survival of IPF is 3.8 years after diagnosis, with lung transplant often the only remaining therapeutic option. Currently approved antifibrotic therapies remain limited in their ability to halt disease progression or reverse established fibrosis, thus presenting an urgent unmet need for further therapeutic development.

Monocyte-derived macrophages (MDMs) are central mediators of pulmonary fibrosis progression. In response to extracellular signals, macrophages dynamically reprogram their metabolic pathways to support specific differentiation states (5–7). While the role of biochemical signals, such as cytokines, in regulating macrophage metabolism has been extensively studied, the mechanisms by which macrophages respond to biophysical cues, such as stiffness, stretch, and tension, are less well understood.

Increased extracellular matrix (ECM) stiffness is a hallmark of fibrotic lung diseases. Whereas the plastic surface of standard tissue culture plates has an approximate stiffness of 10^6^ kPa, the normal lung is highly compliant, with stiffness ranging from 0 to 1 kPa. In contrast, as pulmonary fibrosis progresses, the mechanical properties of aberrant fibrotic tissue change significantly, often reaching stiffness values of 15 to 40 kPa, as measured by atomic force microscopy (AFM). Despite recent advances in understanding mechanosignaling in the fibroblast population and its role in IPF pathogenesis (8, 9), there is limited information on how the mechanical properties of the fibrotic microenvironment alter macrophage phenotypes. A stiffness of ∼150 kPa can increase production of the proinflammatory cytokine TNF-α in macrophages after LPS stimulation (10); however, the mechanism(s) by which macrophages metabolically and functionally adapt to the stiffness levels characteristic of fibrotic lungs remains poorly defined.

Profibrotic MDMs are known to be apoptosis-resistant, and their persistence plays a critical role in fibrosis progression (11–13). While previous studies have shown that mitochondrial reactive oxygen species modulate macrophage apoptosis resistance, little is known about whether physical cues, such as increased ECM stiffness in fibrotic lungs, contribute to MDM apoptosis resistance. While several studies have shown that mechanosignaling can regulate apoptosis in vascular smooth muscle cells and cancer cells (14, 15), none have investigated the effects of ECM stiffness in macrophages.

Anti-apoptotic proteins such as B-cell lymphoma 2 (BCL-2), X-linked inhibitor of apoptosis protein (XIAP), and protein tyrosine phosphatase, non-receptor type 13 (PTPN13) have been previously implicated in IPF. Targeting these anti-apoptotic proteins in fibroblasts can induce fibroblast apoptosis and attenuate or even reverse established pulmonary fibrosis (8, 16–19). Moreover, BCL-2 is upregulated in IPF macrophages (11). However, the role of B-cell lymphoma-extra large (Bcl-xL, encoded by gene BCL2L1), another BCL-2 family protein, has not been evaluated in IPF. BCL-2 and Bcl-xL can differentially regulate apoptosis in immune cells (20), whereas Bcl-xL is more abundant in monocyte-derived macrophages than BCL-2 (21–23).

Here, we demonstrate that mechanoactivated macrophages exhibit apoptosis-resistance, upregulated Bcl-xL expression, and enhanced mitochondrial bioenergetics with increased oxidative phosphorylation (OXPHOS). Mechanoactivated macrophages undergo metabolic reprogramming with augmented glutaminolysis. Inhibiting glutaminolysis enhances apoptosis in mechanoactivated MDMs. Furthermore, selective inhibition of Bcl-xL *in vivo* protected mice against experimental pulmonary fibrosis. Lastly, mechanoactivated macrophages are profibrotic and promote fibrogenesis in precision-cut lung slices (PCLS). Our findings demonstrate a dynamic crosstalk of macrophages and ECM within the fibrotic niche, mediated by matrix stiffness-induced metabolic reprogramming. We conclude that acquired apoptosis resistance in macrophages, in response to mechanoactivation, may represent a potential target for therapeutic intervention in IPF and other fibrotic disorders.

## RESULTS

### Stiffness-activated macrophages acquire apoptosis resistance

We used an asbestos-induced pulmonary fibrosis model to determine the stiffness of fibrotic lung tissue. Lung tissue stiffness in control and chrysotile asbestos-treated mice was measured by AFM microindentation. Fibrotic lungs exhibited significantly increased elastic moduli (Young’s moduli, *E*) compared with nonfibrotic lungs **(Figure 1A)**. The average stiffness of fibrotic lungs, measured as elastic moduli, was 35.02 ± 0.44 kPa, nearly 45-fold higher than that of nonfibrotic lungs (0.77 ± 0.10 kPa) **(Figure 1B)**, consistent with values previously reported (24–26).

**Figure 1:**
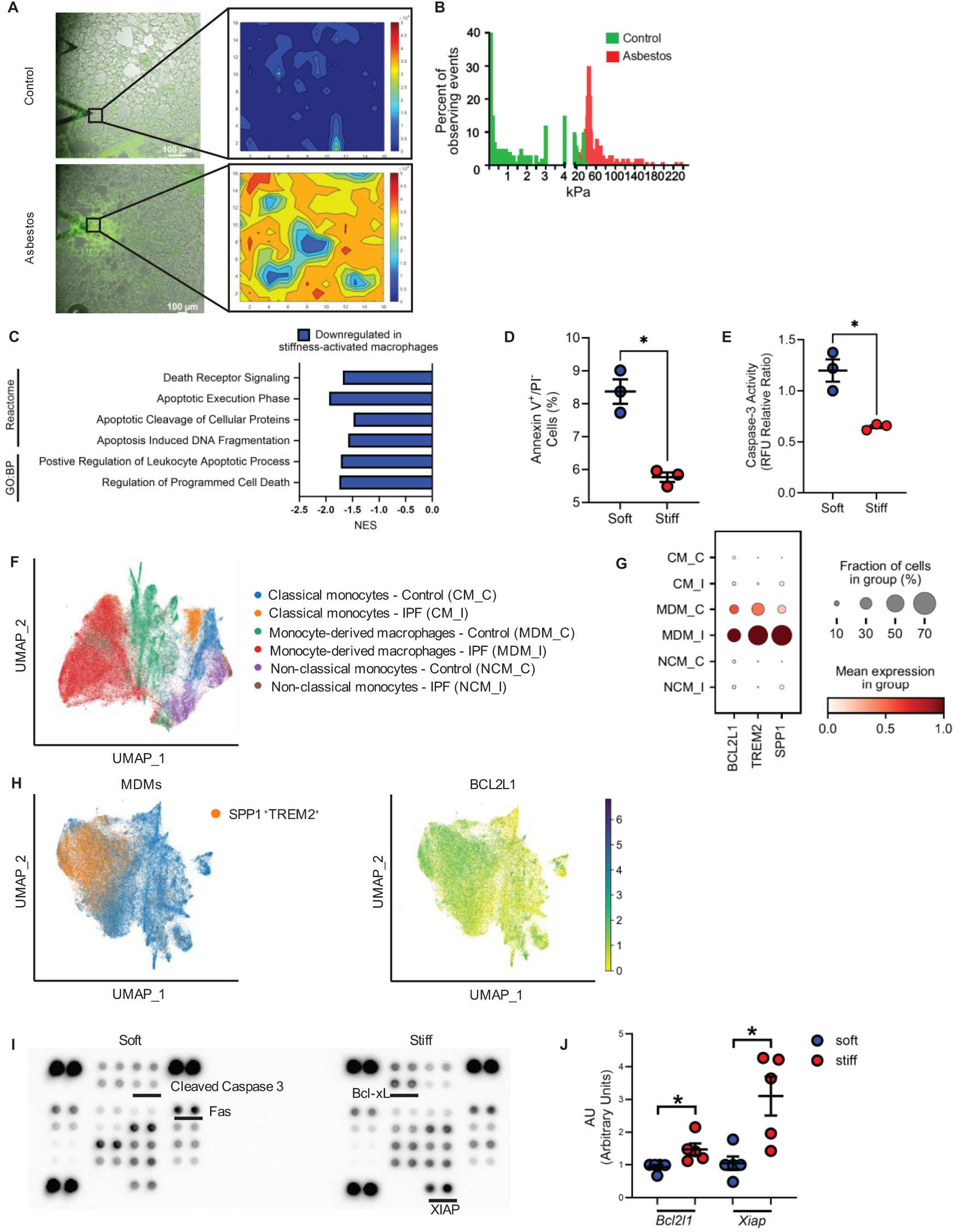
Stiffness-activated macrophages acquire apoptosis-resistance. **(A)** Mice were exposed to chrysotile asbestos (100 μg in 50 μl saline) or control *via* intratracheal instillation on day 1. On day 21, mice were euthanized and frozen lung tissue sections from control- or asbestos-treated mice were stained with green fluorescent Col-F collagen binding reagent to identify areas of fibrosis (*left*). The mechanical properties of the fibrotic lung areas were determined by AFM microindentation. Values of lung tissue stiffness were shown by heatmaps (*right*). Scale bar = 100 μm. **(B)** Histograms showing the elastic moduli distribution in the lungs from control- and asbestos-treated mice. Bone marrow-derived macrophages (BMDMs) were cultured on soft or stiff matrix for 48 hr. **(C)** mRNA was isolated and bulk RNA-seq was performed in BMDMs, and pathway enrichment analysis was performed for Gene Ontology Biological Processes and Reactome pathway compendia. Significantly downregulated pathways in stiffness-activated macrophages are shown. FDR < 0.05. NES: Normalized enrichment score. **(D)** Apoptotic macrophages (Annexin V^+^/PI^-^) were identified by flow cytometry. **(E)** Caspase-3 activity was measured biochemically. **(F)** UMAP plots of scRNA-seq data depicting IPF enriched and Control enriched clusters for lung classical monocytes (CMs), non-classical monocytes (NCM) and MDMs from control (CM_C, MDM_C and NCM_C) (*n* =57) and IPF subjects (CM_I, MDM_I and NCM_I) (*n* = 81) (GSE316367). **(G)** Dot plots showing average expression of *BCL2L1*, *TREM2* and *SPP1* and percentage of cells expressing the gene in lung monocytes and MDMs. **(H)** (left) UMAP plots of MDMs expressing *SPP1* and *TREM2*. (right) Feature plot of scRNA-seq data demonstrating expression of *BCL2L1* in MDMs. **(I)** Apoptosis-related protein levels were measured by using a proteome array. **(J)** *Bcl2l1* and *Xiap* gene expression were measured by RT-PCR. \**P*<0.05. Values are shown as mean ± S.E.M. Two-tailed Welch’s *t*-test was utilized.

Previous studies have shown that IPF lungs exhibit stiffness of >15 kPa (9, 27). We cultured macrophages on polydimethylsiloxane (PDMS) plates with elastic moduli of either 0.5 or 16 kPa (0.5 kPa, hereafter referred to as “soft,” and 16 kPa, hereafter referred to as “stiff”). An ECM Select Array was used to determine the optimal condition for macrophage attachment to the PDMS matrix. We found that vitronectin supported maximal macrophage attachment, whereas native collagen I did not support macrophage attachment to the matrix, as previously reported (28, 29) **(Supplementary Figure 1A and 1 B)**.

Because profibrotic macrophages are apoptosis-resistant, we investigated whether ECM-derived mechanosignaling can modulate macrophage apoptosis. To comprehensively evaluate the transcriptional program in mechanoactivated macrophages, we performed bulk RNA sequencing and gene set enrichment analysis (GSEA) on bone marrow-derived macrophages (BMDMs) from wildtype C57BL/6 mice that were cultured on soft or stiff matrix. GSEA revealed that stiffness-activated macrophages exhibited attenuation of apoptosis-related pathways, based on the Reactome and Gene Ontology Biological Processes pathway compendia **(Figure 1C)**. Consistently, stiffness-activated macrophages displayed reduced Annexin V^+^/PI^-^ staining, suggesting attenuated early apoptosis **(Figure 1D)**. Minimal (<0.05%) late apoptosis (Annexin V^+^/PI^+^) was observed in either soft or stiff-cultured macrophages (data not shown). Furthermore, stiffness-activated macrophages displayed reduced caspase 3 activity, compared to macrophages cultured on soft matrix **(Figure 1E)**.

### Apoptosis-resistance in mechanoactivated macrophages have increased expression of anti-apoptotic proteins

Apoptosis is an active process of programmed cell death regulated by both pro- and anti-apoptotic factors. Known pro-apoptotic factors include Bcl-2-associated X (BAX), BCL2 antagonist/killer 1 (BAK1), and Bcl-2-associated death promoter (BAD), whereas established anti-apoptotic factors include BCL-2, Bcl-xL, myeloid cell leukemia-1 (MCL-1), and XIAP. Using our previously published single-cell RNA sequencing dataset (GSE316367) (30), unbiased clustering using the Leiden approach identified distinct populations of MDM, classical monocyte (CM), and non-classical monocyte (NCM) enriched in IPF and control subjects, respectively, as depicted using uniformed manifold approximation and projection (UMAP) **(Figure 1F)**. We found that *BCL2L1* (the gene encoding Bcl-xL) expression was higher in MDMs from IPF subjects **(Figure 1G)**. IPF MDMs are known to express *SPP1* and *TREM2* (31–34). Next, to determine the expression of *BCL2L1* in MDMs, we performed a focused analysis of the MDMs and annotated the populations that express both *SPP1* and *TREM2* **(Figure 1H, *left*)**. We found enrichment of *BCL2L1*-expressing MDMs in *SPP1^+^TREM2^+^* MDMs **(Figure 1H, *right*)**. To examine if the acquired apoptosis resistance in mechanoactivated macrophages is associated with increased expression of BCL2L1, we cultured BMDMs on soft and stiff matrix and found that stiffness-activated macrophages exhibited increased Bcl-xL and XIAP protein **(Figure 1I)**. Similarly, gene expression levels of *Bcl2l1* and *Xiap* were increased in macrophages cultured on stiff matrix compared with those cultured on soft matrix **(Figure 1J)**.

Taken together, these data demonstrate that mechanosignaling can modulate macrophage apoptosis and that this process correlates with increased expression of anti-apoptotic proteins such as Bcl-xL.

### Stiffness-activated macrophages have enhanced mitochondrial energetics

Metabolic reprogramming is a key process in macrophage polarization and is critical for apoptosis resistance (11–13). We previously demonstrated that sustained OXPHOS is required to maintain the profibrotic phenotype of macrophages (35). However, the upstream signals that drive enhanced OXPHOS remain elusive and no studies have linked mechanosignaling to OXPHOS in profibrotic macrophage polarization.

GSEA revealed that OXPHOS was the most upregulated metabolic hallmark in stiffness-activated macrophages compared with macrophages cultured on soft matrix. Moreover, glycolysis, which is attenuated during profibrotic macrophage polarization, was downregulated in stiffness-activated macrophages **(Figure 2A and 2B)**. Similarly, gene ontology (GO) analyses of biological processes and cellular components indicated that OXPHOS was highly upregulated in stiffness-activated macrophages **(Supplementary Figure 2A and 2B)**. These data suggested that stiffness-activated macrophages exhibit a transcriptional program favoring increased mitochondrial OXPHOS.

**Figure 2:**
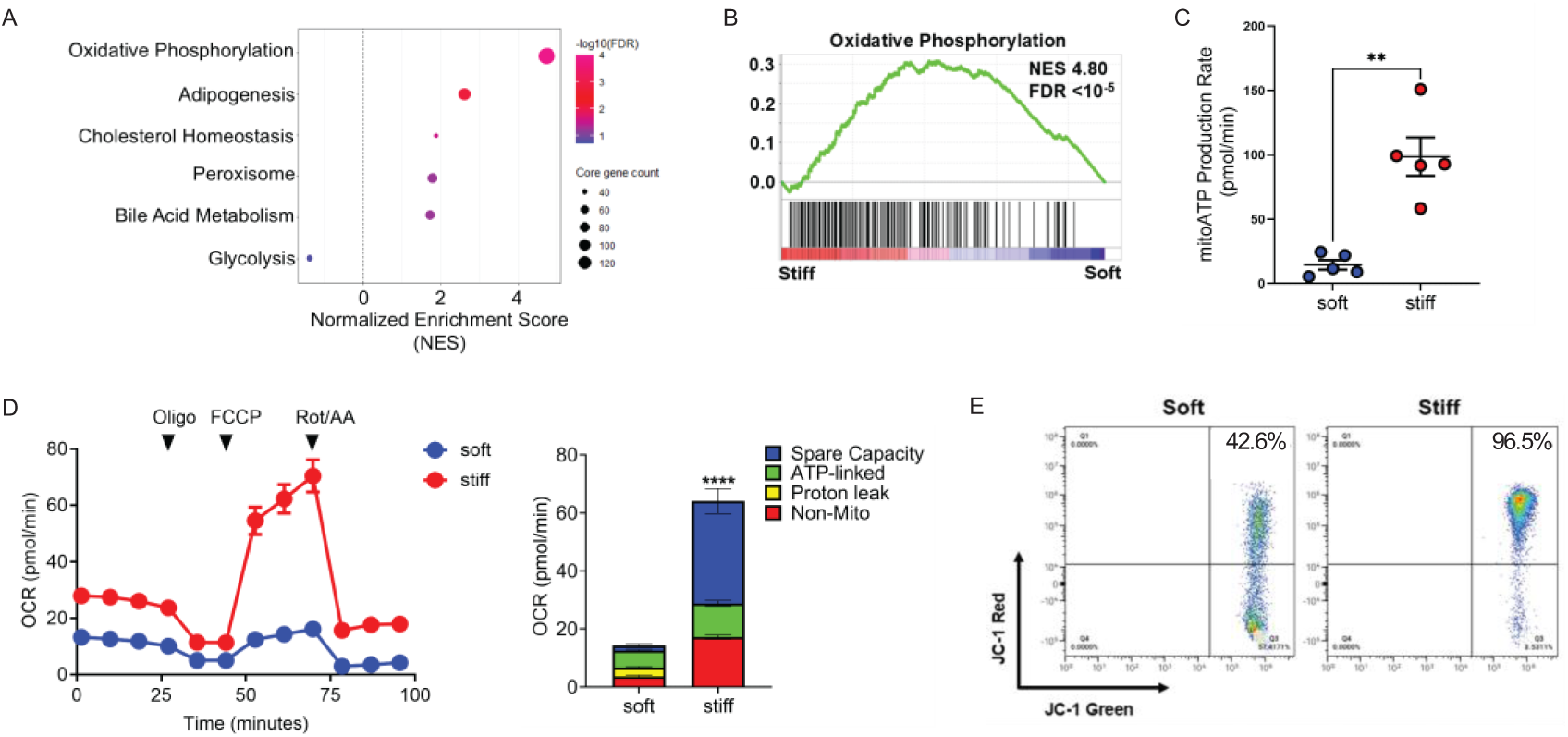
Stiffness-activated macrophages have enhanced mitochondrial energetics. BMDMs were cultured on soft or stiff matrix for 48 hr. **(A)** mRNA was isolated and profiled using bulk RNA-seq, and gene set enrichment analysis (GSEA) was performed. Significantly enriched metabolism-related Hallmark pathways were shown. **(B)** GSEA enrichment plot for the Hallmark OXPHOS pathway. **(C)** Real-time ATP rate assay to measure mitochondrial ATP using Seahorse XFe96. Macrophages were treated sequentially with oligomycin, and Rotenone/Antimycin A (Rot/AA), and the mitoATP production rate was shown. **(D)** Mitochondrial stress test to measure oxygen consumption rate (OCR) using Seahorse XFe96. Macrophages were treated sequentially with oligomycin (Oligo), FCCP, and Rotenone/Antimycin A (Rot/AA) (left). Stacked bar graphs quantify mitochondrial respiration parameters (right). **(E)** Mitochondrial membrane potential was determined by measuring JC-1 dimer/monomer by flow cytometry. \*\**P*< 0.01, \*\*\*\**P*<0.0001. Values are shown as mean ± S.E.M. Two-tailed Welch’s *t*-test was utilized. NES: normalized enrichment score. FDR: false discovery rate

To validate that mechanoactivated macrophages have increased mitochondrial bioenergetics, we first determined whether stiffness could increase mitochondrial ATP production. Oxygen consumption rate (OCR) and real-time ATP production were measured in macrophages cultured on soft and stiff matrix. Stiffness-activated macrophages showed increased mitochondrial ATP production compared with macrophages cultured on soft matrix **(Figure 2C)**. To further delineate mitochondrial bioenergetics, we performed the mitochondrial stress test and found that stiffness-activated macrophages exhibited much higher maximal OCR compared with macrophages cultured on soft matrix, with significant increases in ATP-linked and spare capacity OCR **(Figure 2D)**. Because mitochondrial ATP production is closely related to mitochondrial membrane potential (MMP), we next interrogated whether stiffness-activated macrophages also exhibited increased MMP. We found that macrophages cultured on stiff matrix had higher MMP than macrophages cultured on soft matrix **(Figure 2E)**. Expression of *TFAM*, a key regulator of mitochondrial biogenesis, that is upregulated in fibrotic MDMs (35), was also increased in stiffness-activated macrophages **(Supplementary Figure 2C)**. Taken together, these data demonstrate enhanced mitochondrial bioenergetics in mechanoactivated macrophages.

### Stiffness-induced OXPHOS is fueled by enhanced glutaminolysis

Mitochondrial bioenergetics are supported by three primary substrates: long-chain fatty acids, glucose/pyruvate, and glutamine. Having identified increased mitochondrial bioenergetics in stiffness-activated macrophages, we next sought to determine which metabolic pathway(s) support this enhanced mitochondrial activity. We found that glutaminolysis was the predominant pathway driving the enhanced OCR measured in mechanoactivated macrophages, compared to fatty acid oxidation (FAO) and glycolysis. Inhibition of glutaminolysis with the glutaminase inhibitor, bis-2-(5-phenylacetamido-1,2,4-thiadiazol-2-yl)ethyl sulfide (BPTES) completely reversed the augmented OCR observed in mechanoactivated macrophages, reducing OCR to levels near those found in macrophages cultured on soft matrix **(Figure 3A)**, whereas FAO inhibitor Etomoxir and glycolysis inhibitor UK5099 had minimal effects on OCR in mechanoactivated macrophages.

**Figure 3:**
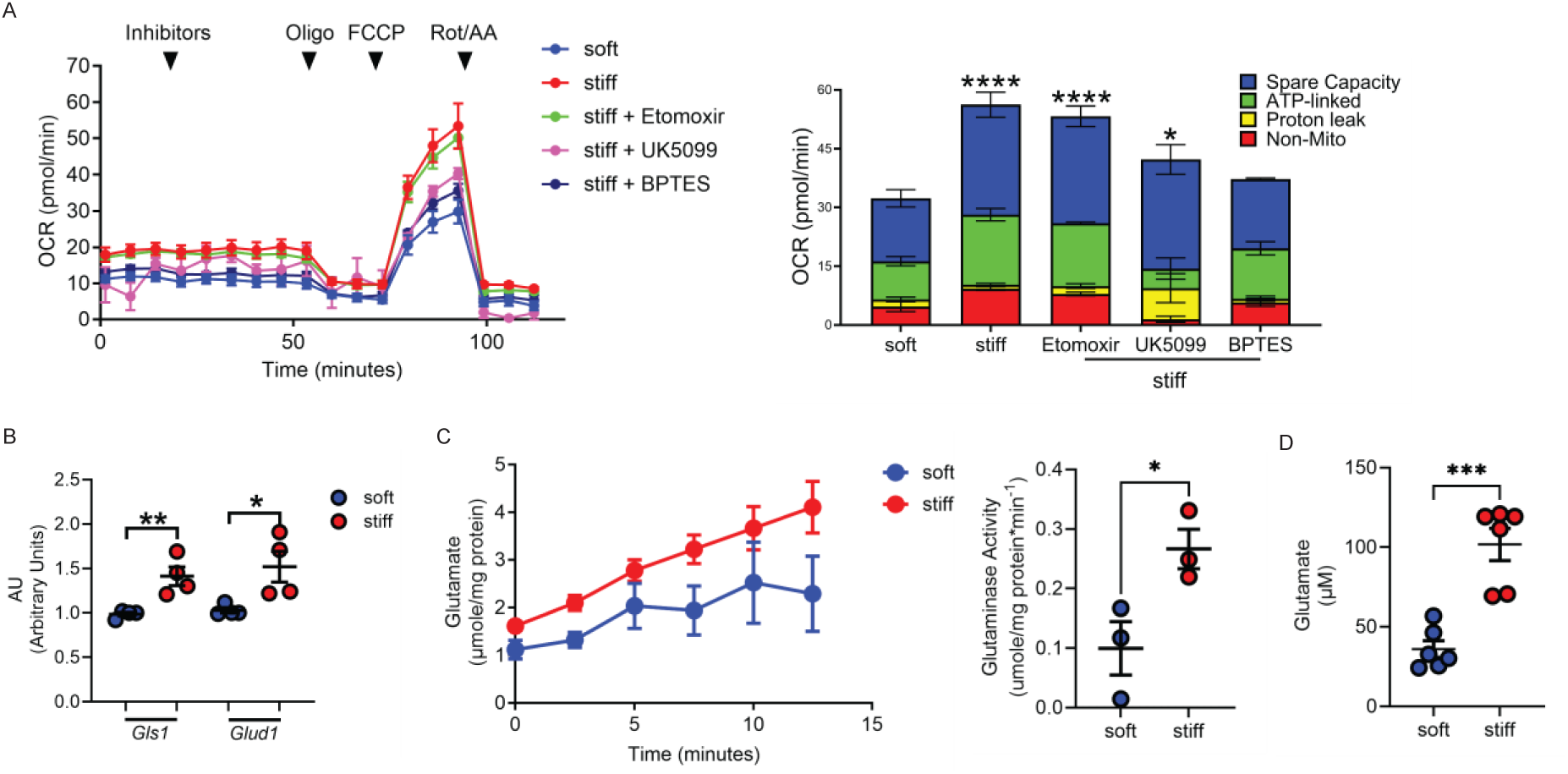
Stiffness-induced OXPHOS is fueled by enhanced glutaminolysis. BMDMs were cultured on soft or stiff matrix for 48 hr. **(A)** Seahorse XF substrate oxidation stress tests were conducted to measure the oxidation of three substrates, long-chain fatty acid (LCFA), glucose/pyruvate, and glutamine, that fuel mitochondrial bioenergetics. Macrophages were treated sequentially with inhibitors (Etomoxir for LCFA, UK5099 for glucose/pyruvate, and BPTES for glutamine), oligomycin, FCCP, and Rotenone/Antimycin A (Rot/AA) (left). Stacked bar graphs quantify mitochondrial respiration parameters (right). **(B)** *Gls1* and *Glud1* gene expression was measured by RT-PCR. **(C)** Glutaminase (GLS) activity was quantified by measuring glutamate production kinetically and as a rate. **(D)** Intracellular glutamate levels were measured biochemically. \**P*<0.05; \*\**P*<0.01, \*\*\**P*<0.001. Values are shown as mean ± S.E.M. Two-tailed Welch’s *t*-test was utilized.

Glutaminolysis is the process by which glutamine is converted to glutamate by glutaminase (GLS) and subsequently to the tricarboxylic acid (TCA) cycle intermediate α-ketoglutarate (α-KG) by glutamate dehydrogenase (GLUD). We found that expression of *Gls1* and *Glud1* were increased in macrophages cultured on stiff matrix compared with macrophages cultured on soft matrix **(Figure 3B)**. We next sought to determine if stiffness could modulate the enzymatic activity of glutaminase and found that glutaminase activity was increased in stiffness-activated macrophages **(Figure 3C)**. Consistently, intracellular glutamate concentrations were also increased in stiffness-activated macrophages **(Figure 3D)**, while glutamine concentrations were reduced **(Supplementary Figure 3A)**. Collectively, these data show that stiffness-activated macrophages exhibit augmented glutaminolysis and establish that this process contributes substantially to the enhanced mitochondrial bioenergetics observed in these cells.

### Enhanced glutaminolysis regulates apoptosis resistance in macrophages

Glutaminolysis is a key mechanism to suppress apoptosis in cancer cells and has been previously implicated in fibroblast activation in IPF (36–38). However, no studies have directly linked mechanoactivation to glutaminolysis and apoptosis regulation in profibrotic macrophages. To determine whether glutaminolysis regulates macrophage apoptosis resistance, macrophages were cultured on either soft or stiff matrix in the presence or absence of the GLS inhibitor BPTES. We found that BPTES-treated macrophages exhibited increased apoptosis, even when cultured on stiff matrix. In contrast, addition of dimethyl-glutamate (DM-Glu), a cell-membrane-permeable glutamate analog (39), bypassed BPTES-mediated GLS inhibition and rescued macrophages from BPTES-mediated apoptosis **(Figure 4A)**.

**Figure 4:**
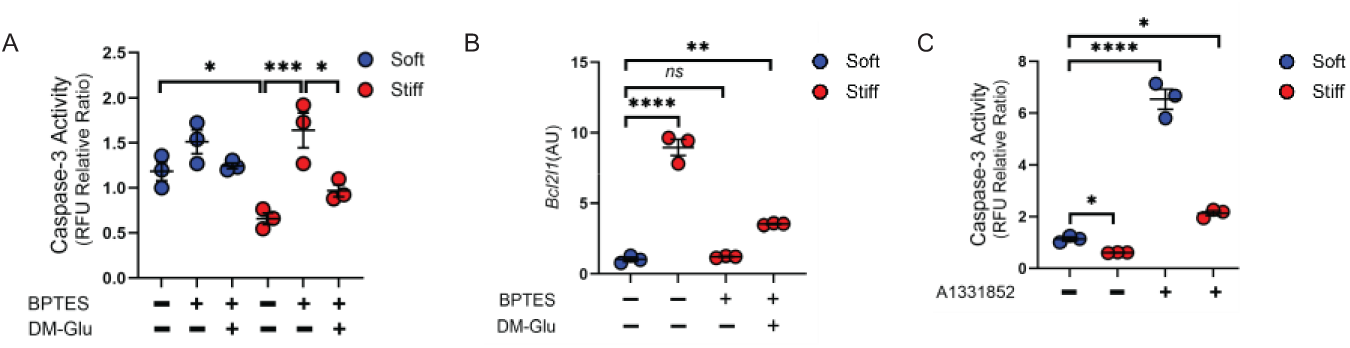
Enhanced glutaminolysis regulates apoptosis resistance in macrophages. **(A)** BMDMs were cultured on soft or stiff matrix in the presence or absence of the GLS inhibitor BPTES (3 μM) and DM-Glu (5 mM) for 48 hr. Caspase-3 activity was measured. **(B)** *Bcl2l1* expression was measured by RT-PCR. **(C)** BMDMs were cultured on soft or stiff matrix for 48 hr and then treated with either vehicle or A-1331852 for 4 hr. Caspase-3 activity was measured. \**P*<0.05; \*\**P*<0.01, \*\*\**P*<0.001, \*\*\*\**P*<0.0001. Values are shown as mean ± S.E.M. One-way ANOVA followed by Tukey’s multiple comparison test was utilized. DM-Glu: Dimethyl glutamate

To determine whether Bcl-xL expression depends on glutaminolysis, macrophages were cultured on either soft or stiff matrix in the presence or absence of BPTES. We found that *Bcl2l1* expression was reduced in BPTES-treated macrophages, even when cultured on stiff matrix. In contrast, the addition of DM-Glu rescued BPTES-mediated suppression of *Bcl2l1* expression **(Figure 4B)**. Similarly, BPTES attenuated *Xiap* expression in stiffness-activated macrophages, whereas the addition of DM-Glu reversed this inhibition **(Supplementary Figure 4A).**

To determine whether Bcl-xL inhibition increases apoptosis in stiffness-activated macrophages, macrophages were cultured in the presence or absence of A-1331852, a selective Bcl-xL inhibitor with proven efficacy in cancer models (40, 41). We found that A-1331852 induced apoptosis in macrophages, even in those cultured on stiff matrix **(Figure 4C)**. Taken together, these findings indicate that enhanced glutaminolysis contributes to stiffness-mediated macrophage apoptosis resistance, at least in part, through upregulation of Bcl-xL.

### Inhibiting Bcl-xL attenuates pulmonary fibrosis *in vivo*

Apoptosis-related pathways have been explored as therapeutic targets in pulmonary fibrosis. Targeting Bcl-2 with the selective inhibitor ABT-199 (venetoclax) can protect mice from developing pulmonary fibrosis, reverse established fibrosis, and promote fibrosis resolution. The relevant cellular targets of Bcl-2 inhibition may include profibrotic macrophages (11) or myofibroblasts (16), as both populations acquire apoptosis resistance in pulmonary fibrosis. However, the effects of specifically targeting Bcl-xL have not been evaluated *in vivo*.

To determine the effect of selectively targeting Bcl-xL *in vivo*, wild-type mice were exposed to chrysotile asbestos or control on day 1. Beginning on day 10, after fibrosis had initiated (42), mice received vehicle control or the selective Bcl-xL inhibitor, A-1331852, (25 mg/kg) twice daily by oral gavage until day 21, when fibrosis endpoints were assessed **(Figure 5A)**. Mice treated with A-1331852 showed reduced lung collagen deposition on histological evaluation, whereas vehicle-treated mice exhibited architectural distortion and aberrant collagen accumulation on day 21 **(Figure 5B)**. These histologic findings were further confirmed biochemically by hydroxyproline analysis. The asbestos-injured mice treated with A-1331852 had reduced lung hydroxyproline compared to asbestos-injured mice that received the vehicle **(Figure 5C)**. In lung tissue, A-1331852 significantly increased the number of TUNEL^+^ cells, compared to mice that received the vehicle **(Figure 5D-E)**. In addition, mice treated with A-1331852 had lower levels of TGF-β1 **(Figure 5F)** and secreted phosphoprotein 1 (SPP1) **(Figure 5G)** in bronchoalveolar lavage fluid after chrysotile asbestos exposure compared with vehicle-treated mice. Taken together, these data suggest that pharmacologic targeting of Bcl-xL protects mice from pulmonary fibrosis even after fibrotic remodeling has been initiated.

**Figure 5:**
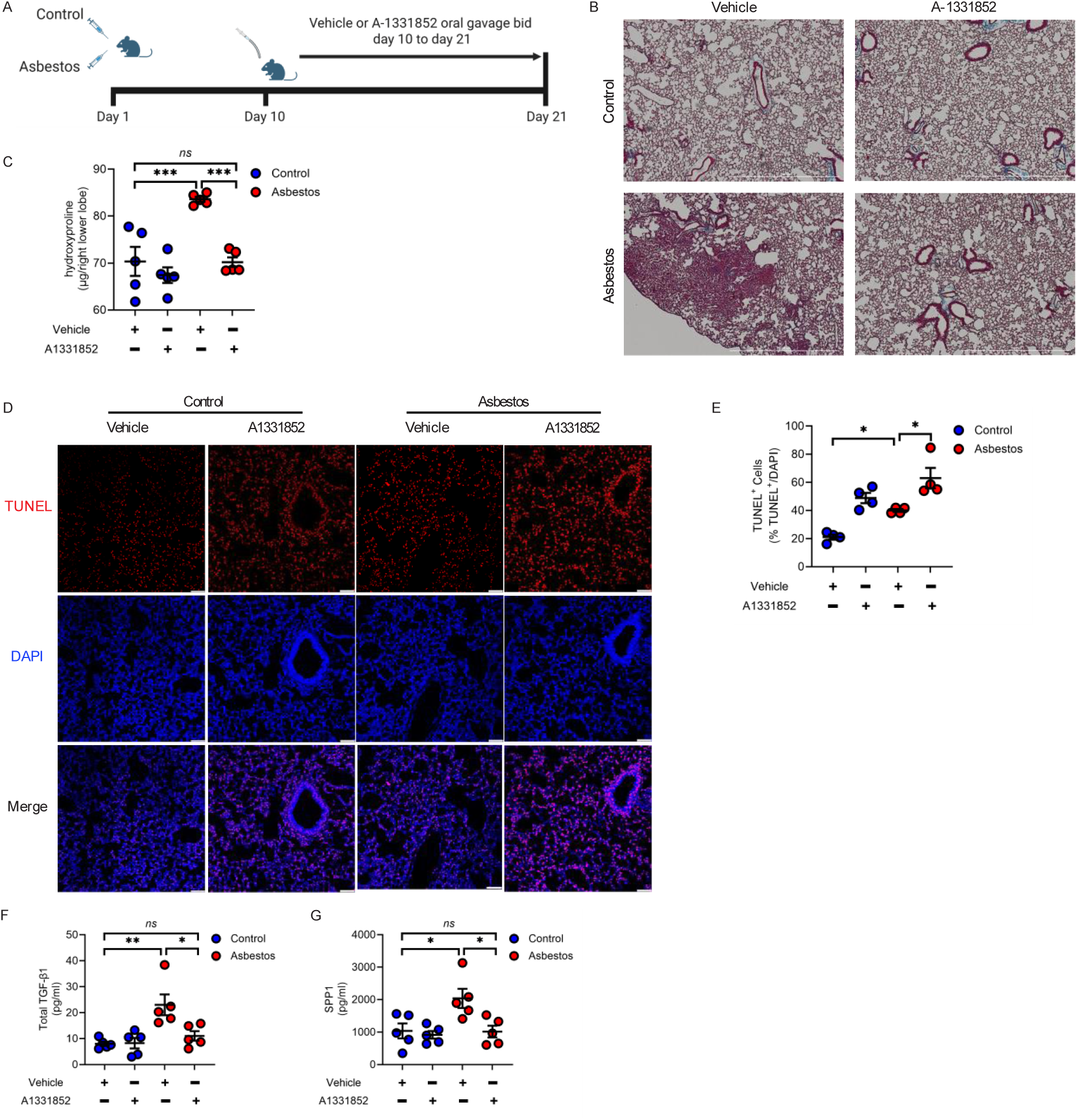
Wild type mice were exposed to either saline or chrysotile asbestos on day 1. Beginning on day 10, mice received vehicle control or A-1331852 (25 mg/kg) twice daily by oral gavage until day 21, when they were euthanized, and lung tissues were harvested. **(A)** Schematic of the i.t. exposure, A-1331852 treatment, and harvesting schedule. **(B)** Representative histology of lung sections (*n* = 5 per group). Bar indicates 1000 μm. **(C)** Hydroxyproline analysis of homogenized lung (*n* = 5 per group). **(D)** Representative images of lung IF for TUNEL (red) and DAPI (blue). Scale bars: 50 μm. **(E)** Quantification of TUNEL^+^ cells. **(F)** Total TGF-β1 level and **(G)** SPP1 level in BAL fluid measured by ELISA. \**P*<0.05; \*\**P*<0.01, \*\*\**P*<0.001. Values are shown as mean ± S.E.M. One-way ANOVA followed by Tukey’s multiple comparison test was utilized. Each dot represents one animal.

### Mechanoactivated macrophages are profibrotic and promote ECM production

Profibrotic MDMs are known to generate growth factors such as TGF-β1 (12) to stimulate fibroblasts, the principal effector cells responsible for ECM production. We found that stiffness-activated primary lung macrophages produced higher amounts of TGF-β1 and SPP1 than macrophages cultured on the soft matrix **(Figure 6A and 6B)**. Moreover, we found that while stiffness promoted TGF-β1 production in mechanoactivated macrophages, BPTES attenuated TGF-β1 production in these cells. In contrast, DM-Glu restored TGF-β1 production, even in the presence of BPTES **(Figure 6C)**.

**Figure 6:**
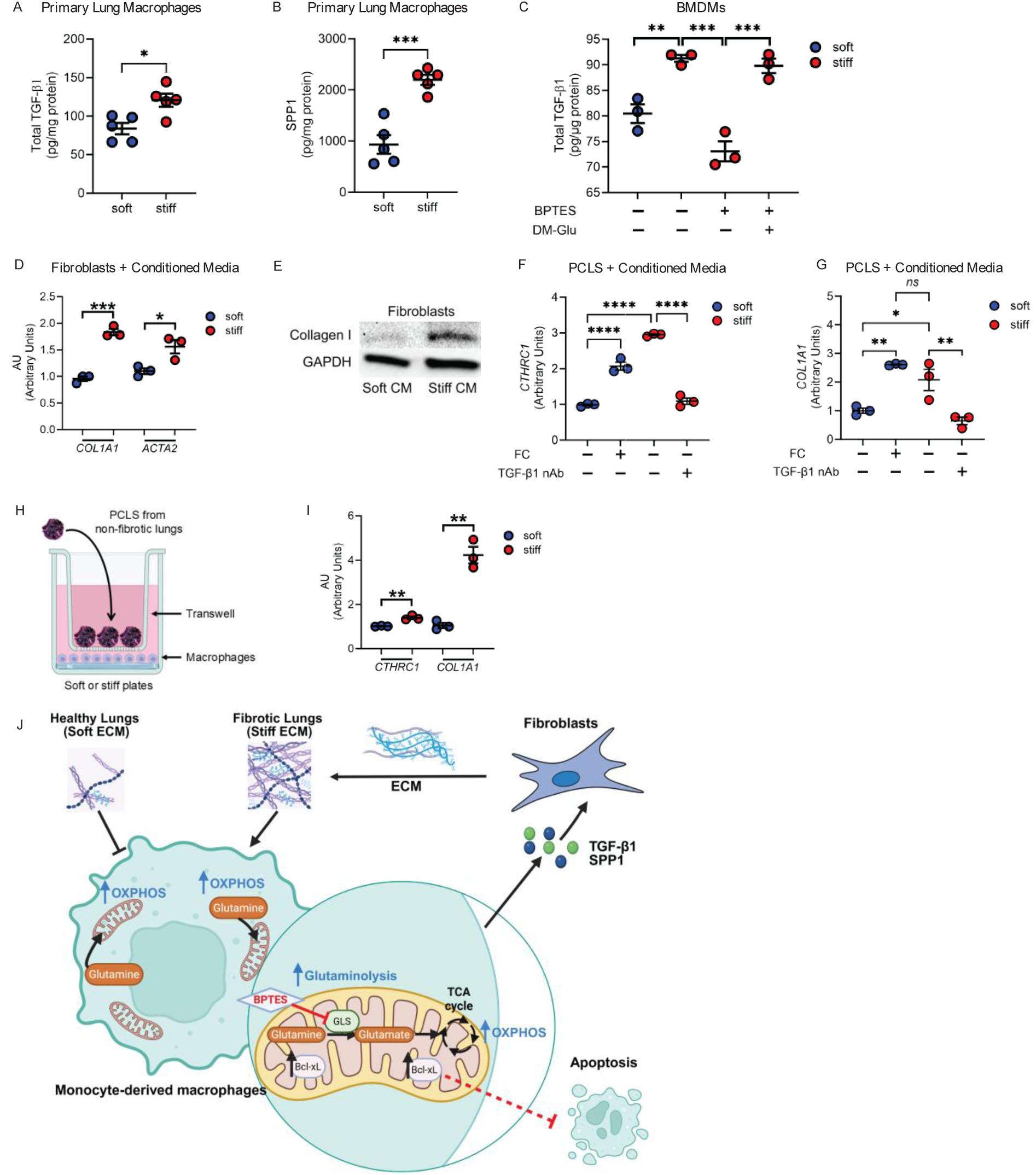
Mechanoactivated macrophages are profibrotic and promote ECM production. Primary lung macrophages were isolated from IPF lungs and cultured on soft or stiff matrix for 48 hr. **(A)** TGF-β1 level and **(B)** SPP1 levels in the supernatant were measured by ELISA. *n* =4. BMDMs were cultured on soft or stiff matrix in the presence or absence of the GLS inhibitor BPTES (3 μM) and DM-Glu (5 mM) for 48 hr. **(C)** TGF-β1 level in the supernatant were measured by ELISA. Primary lung macrophages were isolated from IPF lungs and cultured on soft or stiff matrix for 48 hr. Conditioned media were subsequently collected. **(D)** Primary lung fibroblasts were obtained from non-fibrotic lungs and cultured with the conditioned media for 48 hr. mRNA was isolated from the fibroblasts. *COL1A1* and *ACTA2* mRNA expression were measured by RT-PCR. **(E)** Fibroblast cell lysates were used for immunoblot analysis, and Collagen I protein were measured. Precision-cut lung slices (PCLS) were obtained from non-fibrotic lungs and cultured with the conditioned media in the presence or absence of fibrotic cocktail (FC) or anti-human TGF-β1 neutralizing antibody (0.125 μg/ml) for 48 hr. mRNA was isolated from the PCLS. **(F)** *CTHRC1* and **(G)** *COL1A1* mRNA expression levels were measured by RT-PCR. *n*=3. **(H)** Primary lung macrophages were isolated from IPF lungs and cultured on soft or stiff matrix. PCLS generated from non-fibrotic lungs were cultured in Transwell inserts placed above the primary lung macrophages. The macrophages and PCLS were co-cultured for 48 hr. **(I)** mRNA was isolated from the PCLS. *CTHRC1* and *COL1A1* mRNA expression were measured by RT-PCR. *n* =3. **(J)** Schematic depicting how lung tissue stiffness promotes macrophage apoptosis resistance *via* metabolic reprogramming in pulmonary fibrosis. \**P*<0.05; \*\**P*<0.01, \*\*\**P*<0.001, \*\*\*\**P*<0.0001. Values are shown as mean ± S.E.M. Two-tailed Welch’s *t*-test or one-way ANOVA followed by Tukey’s multiple comparison test was utilized.

To determine the profibrotic effects of mechanoactivated macrophages and their intercellular crosstalk with other cells critical for fibrosis development, we cultured primary IPF macrophages on soft and stiff matrix and collected conditioned media. We then cultured primary human lung fibroblasts derived from non-fibrotic subjects with the conditioned media. We found that conditioned media from stiffness-activated macrophages increased *COL1A1* and *ACTA2* expression in lung fibroblasts **(Figure 6D)**. Similarly, there was increased collagen production in lung fibroblasts cultured with conditioned media from stiffness-activated macrophages **(Figure 6E)**.

To determine whether mechanoactivated macrophages can induce profibrotic responses within the complex multicellular environment of human lung tissue, we next performed studies using PCLS. PCLS are *ex vivo* tissue sections that preserve native architecture and cellular diversity, providing a physiologically relevant model to study immune and fibrotic responses under controlled conditions (43). PCLS derived from non-fibrotic human lungs were cultured with conditioned media from IPF macrophages. We found that PCLS cultured with conditioned media from stiffness-activated IPF macrophages exhibited increased *CTHRC1* and *COL1A1* expression, compared to PCLS cultured with conditioned media from IPF macrophages cultured on soft matrix or PCLS treated with the fibrotic cocktail (44) **(Figure 6F and 6G)**. More importantly, the addition of anti-TGF-b1 neutralizing antibody in the conditioned media abrogated the augmented *CTHRC1* and *COL1A1* expression, suggesting that stiffness-activated macrophages promote fibrosis *via* profibrotic cytokines such as TGF-β1.

Lastly, we performed co-culture experiments using PCLS derived from non-fibrotic lungs and mechanoactivated primary lung macrophages from IPF subjects **(Figure 6H)**. PCLS derived from non-fibrotic subjects co-cultured with stiffness-activated macrophages displayed increased expression of *CTHRC1* and *ACTA2* **(Figure 6I)**. Taken together, these data suggest that mechanoactivated macrophages produce profibrotic cytokines such as TGF-β1, and enhance ECM production, and thereby promote fibrosis progression.

## DISCUSSION

Monocyte-derived macrophages are increasingly recognized as key mediators of IPF progression. Although increased ECM stiffness is a defining feature of fibrotic lung disease, the mechanisms by which macrophages adapt to the mechanical properties of the fibrotic niche remain incompletely understood. A notable finding of this study is that mechanical cues from the stiffened extracellular environment are sufficient to promote metabolic reprogramming in macrophages. We show that mechanoactivated macrophages acquire an apoptosis-resistant profibrotic phenotype and exhibit enhanced mitochondrial bioenergetics. Importantly, this metabolic reprogramming is driven predominantly by augmented glutaminolysis. Inhibition of glutaminolysis increased apoptosis in mechanoactivated MDMs, suggesting that metabolic reprogramming is a key determinant of profibrotic macrophage apoptosis-resistance in pulmonary fibrosis **(Figure 6J)**. By showing that matrix stiffness enhances glutaminolysis, sustains mitochondrial OXPHOS, and promotes Bcl-xL-dependent apoptosis resistance, this study provides a framework for understanding how physical cues in the fibrotic niche shape macrophage fate and function in IPF.

Glutaminolysis is a critical metabolic process which has previously been implicated in pulmonary fibrosis, particularly in fibroblasts. Previous studies have shown that *Gls1* expression and glutamine metabolism are increased in bleomycin-injured mouse lungs and that TGF-β1 treatment induces *Gls1* expression in fibroblasts (45, 46). In TGF-β1-stimulated fibroblasts, increased GLS1 promotes collagen translation and limits its degradation, thereby promoting ECM accumulation, and augmented glutaminolysis is crucial for myofibroblast differentiation (47). Genetic silencing of *Gls1* in fibroblasts or therapeutic targeting of *Gls1* with a chemical inhibitor *in vivo* can protect mice from developing pulmonary fibrosis (45). In addition, glutaminolysis suppresses fibroblast apoptosis, which may further contribute to fibrosis progression (48). Interestingly, inhibition of glutaminolysis in fibroblasts has no effect on mitochondrial bioenergetics (49). Our findings extend the paradigm of metabolic reprogramming towards glutaminolysis in IPF beyond the fibroblast population and identify macrophages as an additional cell population in which augmented glutaminolysis contributes to fibrosis development. More importantly, our study demonstrates that mechanosignaling can directly modulate metabolic reprogramming in MDMs, a previously unidentified mechanism.

Apoptosis resistance is a key feature of progressive pulmonary fibrosis, enabling the persistence of pathogenic mesenchymal and immune cell populations that drive ECM deposition and tissue remodeling. Fibroblasts in fibrotic lung tissue exhibit enhanced survival signaling and resistance to apoptosis, contributing to aberrant wound healing and fibrosis progression (16–19). In this study, we establish a novel role for the anti-apoptotic protein Bcl-xL in the maintenance of profibrotic macrophages. We found that stiffness can reprogram macrophage survival pathways by increasing Bcl-xL expression. Rescue of BPTES-induced apoptosis by DM-Glu further supports the conclusion that glutamine-derived metabolites are functionally linked to macrophage survival under stiff conditions. Moreover, direct pharmacologic inhibition of Bcl-xL abolished stiffness-mediated apoptosis resistance, placing Bcl-xL downstream of this mechanometabolic program. These findings suggest that increased matrix stiffness actively reinforces macrophage persistence by coupling metabolic and survival signaling. In the fibrotic lung, where tissue stiffening intensifies with disease progression, this mechanism may create a feed-forward loop in which stiffened ECM sustains long-lived, glutaminolysis-dependent mechanoactivated macrophages, which subsequently secrete profibrotic cytokines. Thus, targeting a mechanoactivated glutaminolysis-Bcl-xL axis may disrupt macrophage persistence and limit fibrosis progression.

Targeting anti-apoptotic signaling using BH3 mimetics has emerged as a promising therapeutic strategy. The pan-BCL-2 family inhibitor, ABT-263, can target myofibroblasts and reverse established pulmonary fibrosis *in vivo* and bleomycin-induced skin fibrosis (17, 50). ABT-199 (also known as venetoclax), a selective BCL-2 inhibitor approved for treating several forms of leukemia (51), attenuated pulmonary fibrosis in independent preclinical studies by targeting fibroblasts and macrophages separately (11, 16). BCL-2 and Bcl-xL can differentially regulate apoptosis in immune cells (20), whereas Bcl-xL is more abundant in monocyte-derived macrophages than BCL-2 (21–23). Antibody-conjugated A-1331852, mirzotamab clezutoclax (ABBV155), has shown efficacy in non-human primates and a safety profile in phase 1b clinical trials (52, 53). We also found that *Tfam* expression is increased in stiffness-activated macrophages. TFAM has been shown to modulate BCL2L1 expression in ovarian cancer cells (54). In addition to MDMs, Bcl-xL is also increased in α-SMA^+^ fibroblasts from IPF and silicosis subjects (17) and is expressed in alveolar type II cells (55). Besides Bcl-xL, we also found that mechanoactivated macrophages displayed increased expression of XIAP, an anti-apoptotic protein known to mediate apoptosis in macrophages (56, 57). The role of XIAP in mechanoactivated macrophages and its therapeutic implication warrants further investigation. In summary, our study on Bcl-xL, together with other studies (11, 16, 17), suggests that restoring apoptotic sensitivity in profibrotic and apoptosis-resistant cell populations may represent an effective therapeutic strategy in pulmonary fibrosis.

Collectively, our findings identify a mechanometabolic pathway through which the stiff fibrotic microenvironment promotes macrophage metabolic reprogramming and resistance to apoptosis. Our data suggest that interrupting macrophage survival pathways may help deplete persistent profibrotic macrophages. Since glutaminolysis has also been implicated in fibroblast activation, targeting this pathway may provide the additional advantage of simultaneously modulating multiple profibrotic cell populations. More broadly, our results highlight the importance of integrating mechanobiology with immunometabolism in the study of pulmonary fibrosis. We suggest that metabolic reprogramming of macrophages and associated apoptosis resistance, in response to mechanosignaling in fibrotic tissue, may represent a promising avenue for therapeutic intervention in fibrotic disease.

## METHODS

### Sex as a biological variable

For human tissue-based studies, both male and female individuals were used. For mouse studies, we examined male and female animals, and similar findings are reported for both sexes. Data presented in this manuscript includes both sexes.

### Cell culture and reagents

Mouse BMDMs were generated as previously described (58). Briefly, bone marrow cells from C57BL/6 mice were cultured in M-CSF-supplemented (20 ng/mL) media for 7 days, followed by F4/80 selection (StemCell Technologies, Vancouver, CA). Primary lung macrophages were isolated from explanted lungs from IPF subjects. Briefly, IPF lungs were digested to obtain a single cell suspension. Cells were cultured on non-tissue culture-treated plates for 90 minutes and then attached cells were subjected to CD14 selection (StemCell Technologies, 17858). CD14^+^ cells were cultured on the CytoSoft plates. The ECM Select Array Kit Ultra (5170, Advanced BioMatrix, Carlsbad, CA) was used to select the optimal ECM for cell culture according to the manufacturer’s protocol, as previously described (59, 60). CytoSoft plates coated with polydimethylsiloxane (PDMS) with defined elastic moduli of either 0.5 kPa or 16 kPa were purchased from Advanced BioMatrix.

### Mouse model

Male and female wild type C57BL/6 mice, 8-12 weeks old, were administered chrysotile asbestos (100 μg suspended in 50 μL 0.9% saline solution, UCC Calidria Chrysotile Lot. R-G 144) or titanium dioxide (TiO_2_, 100 μg/50 μl, Sigma Aldrich, 224227), as a control, intratracheally after being anesthetized with 3% isoflurane using a precision Fortec vaporizer (Cyprane). Starting from day 10, A-1331852 (25 mg/kg, AmBeed) was given *via* oral gavage twice a day, as previously described (40). On day 21, mice were euthanized and BAL was performed. The lungs were removed and stained for collagen fibers using Masson’s trichrome stain.

### Mechanical testing of the lung tissue

Stiffness of lungs was mechanically characterized using an MFP-3D-BIO Atomic Force Microscope (Asylum Research), mounted on a Nikon Eclipse Ti2 microscope, in contact mode as previously described (26).

### Bulk RNA sequencing

BMDMs were cultured on soft and stiff matrix for 48 hr. Total RNA was obtained using Quick-RNA Microprep kit (Zymo Research). Total RNA quality was assessed using a NanoDrop spectrophotometer (ThermoFisher Scientific), Qubit 2.0 fluorometric quantitation (Invitrogen), and RNA integrity was evaluated with a 2100 Bioanalyzer and RNA chips (Agilent Technologies, Santa Clara, CA). Libraries were prepared from 200 ng of total RNA using the NEBNext Ultra II Directional RNA-Seq Sample Preparation with rRNA Depletion protocol (Cat. # E7760). The resulting libraries were quantified by Qubit 2.0 fluorometric quantitation and assessed for fragment size distribution using the Bioanalyzer. Equimolar libraries were pooled and sequenced on the Illumina NovaSeq X platform using a 10B flow cell with 2% PhiX spike-in for quality control. Paired-end 150-bp sequencing generated an average of 89 million read pairs per sample.

### Bioinformatics analysis

For bulk RNA-Seq data, adapter sequences and low quality basepairs were trimmed using trim_galore. Data was mapped to the mouse genome build mm39 using STAR v2.7.10b (61). Gene expression was quantified using featureCounts (62) and the Gencode gene model. Differentially expressed genes (DEGs) were determined using EdgeR (63) with significance achieved at FDR<0.05 and fold change exceeding 1.5x. Enriched pathways were determined using the Gene Set Enrichment Analysis (GSEA) method (64), with significance achieved at FDR<0.25 per best practices recommended by the method authors.

Single cell lung data was processed using the Python Scanpy library (65). We used a single human pulmonary fibrosis dataset (30), annotated using CellTypist (66) based on the Human Lung Cell Atlas (HLCA) (67) reference. UMAP plots were generated using Scanpy. We processed separately the monocyte derived macrophages (MDM), classical monocytes (CM), and non-classical monocytes. For each cell type, we performed batch correction using SCVI (68) with the sequencing batch as main covariate; next, unbiased clustering was performed using the Leiden approach, and two clusters were determined per cell type. Dotplots for individual genes were generated using the Scanpy library.

### Flow cytometry-based annexin V measurement

FITC Annexin V Apoptosis Detection Kit with PI (640914, BioLegend) was used to quantify annexin V-positive live cells, according to the manufacturer’s instructions.

### Apoptosis array

The relative levels of apoptosis-related proteins in BMDMs were measured by using a Proteome Profiler Mouse Apoptosis Array (ARY031, R&D Systems), according to the manufacturer’s instructions.

### Caspase 3 activity

Caspase-3 activity was quantitated with the EnzCheck Caspase-3 Activity Assay (Invitrogen E13184) as described in the manufacturer’s instructions.

### Mitochondrial respiration measurement

Oxygen consumption rate (OCR) was determined by using a Seahorse XF96 bioanalyzer (Seahorse Bioscience, Billerica, MA, USA). In brief, BMDMs were cultured on soft and stiff matrix for 48 hr. Macrophages (2.5 x 10^4^) were placed in an XF96 cell culture microplate for 2 hr in DMEM media containing 10 mM glucose, 1 mM pyruvate, and 2 mM glutamine at 37°C without CO_2_. The plate was then subjected to OCR measurement in the XF96 extracellular flux analyzer. For mitochondrial OCR measurement, the following compounds were sequentially added: oligomycin (2 μM), carbonyl cyanide 4-(trifluoromethoxy)phenylhydrazone (FCCP) (4 μM), and antimycin A/rotenone (1 μM). For substrate oxidation measurement, the following inhibitors were added: etomoxir (final dose 4 μM), UK5099 (final dose 2 μM), and BPTES (final dose 3 μM). For real-time ATP rate assay, oligomycin (2 μM) and antimycin A/rotenone (1 μM) were sequentially added.

### Flow cytometry-based MMP measurement

JC-1 mitochondrial membrane potential dye (65-0851, Invitrogen) was used to determine MMP, according to the manufacturer’s instructions.

### RNA extraction and quantitative real-time PCR analysis

Total RNA was obtained using the Quick-RNA Microprep kit (Zymo Research). After reverse transcription using the iScript reverse transcription kit (Bio-Rad Laboratories) was completed, specific gene mRNA expression was determined by quantitative real-time PCR using the SYBR Green kit (Bio-Rad Laboratories). Data were calculated by the ^ΔΔ^CT method. The mRNA measurements were normalized to *Rpl13* and expressed in arbitrary units.

### Glutaminase activity, glutamine, and glutamate concentration measurement

Glutaminase activity was determined by using a QuantiFluo Glutaminase Assay kit (DGLN-100, BioAssay Systems) according to the manufacturer’s instructions. Glutamine concentration was measured using an EnzyChrom Glutamine Assay kit (EGLN-100, BioAssay Systems), and glutamate concentration was measured using an EnzyChrom Glutamate Assay kit (EGLT-100, BioAssay Systems).

### TUNEL assay

Mouse lung sections (5-μm-thick) were prepared, deparaffinized, rehydrated, and permeabilized as previously described (58). Cells were subjected to TUNEL staining using the In-Situ Cell Death Detection Kit, TMR Red (Roche # 12156792910), according to the manufacturer’s instructions. Cells were counterstained with DAPI. TUNEL staining was evaluated by confocal analysis, and data were quantified as TUNEL density over the corresponding DAPI area using ImageJ. Images were captured with a Leica Stellaris confocal microscope.

### Precision cut lung slices (PCLS), conditioned media and co-culture experiments

PCLS from deidentified non-fibrotic control lungs (“failed donor”) from Baylor College of Medicine Chronic Lung Disease Tissue Repository were prepared as previously described (43). Primary lung macrophages were cultured on soft or stiff matrix and PCLS were incubated in Transwell insert (4450, Corning) for 48 h. For the conditioned media experiments, PCLS were cultured in the presence of the fibrotic cocktail, as previously described (44). To neutralize TGF-β1 in the conditioned media, Ultra-LEAF anti-human TGF-β1 recombinant antibody (0.125 mg/ml, 947303, BioLegend) was added. Slices were subjected to total RNA isolation to measure gene expression.

### Hydroxyproline determination

Lung tissue was dried and digested for 24 hours at 112°C with 6N hydrochloric acid. Hydroxyproline concentration was determined as previously described (69).

### Statistics

Statistical analyses were performed using Graphpad Prism 10 statistical software. A minimum of three biological replicates were required in determining the statistical significance. All data were expressed as mean ± SEM. Normal distribution was analyzed to determine whether data met the assumptions of the statistical test. Groups with similar variances were used for statistical comparisons. Statistical analyses were performed with either an unpaired two-sided Welch’s *t*-test, one-way ANOVA with Tukey’s multiple comparison test, or two-way ANOVA. *p* < 0.05 is considered significant.

### Study approval

We obtained non-fibrotic lung fibroblasts, non-fibrotic explanted lungs, explanted lungs with pulmonary fibrosis under approved protocols (H-46823) by the Human Subjects Institutional Review Board of Baylor College of Medicine. All participants provided prior written consent to participate in the study. Animal experiments were approved by the Baylor College of Medicine Institutional Animal Care and Use Committee under protocols AN-9386 and were performed in accordance with the NIH Guide for the Care and Use of Laboratory Animals.

## Data availability

Data are available in the Supporting Data Values file or associated supplemental files. scRNA-seq data are publicly available with accession number of GSE316367. Bulk RNA sequencing data were deposited in the Gene Expression Omnibus (GEO) database with accession numbers GSE344095.

## AUTHOR CONTRIBUTIONS

CH, CC, SKA, FK, YZ, ABC and IOR conceived and designed the study. CH, YZ, ABC and IOR provided the reagents. CH, CC, NG, COL, HG, ERE, XJ, AWC, JAZ, YZ and IOR conducted the experiments and acquired the data. CH and CC wrote the manuscript. LJC, SAO, NJM, JLC, SKA, FK, YZ, ABC and IOR assisted in manuscript editing.

## ACKNOWLEDGMENTS

We appreciate the suggestions from Dr. Stefan Ryter on the manuscript. Chrysotile asbestos was generously provided by Dr. Peter S. Thorne, University of Iowa College of Public Health.

