## supplementary for "Mechanosignaling promotes macrophage apoptosis resistance in pulmonary fibrosis *via* metabolic reprogramming"

**Supplementary Figure 1: Stiffness-activated macrophages are apoptosis resistance.**

Supplementary Figure 1

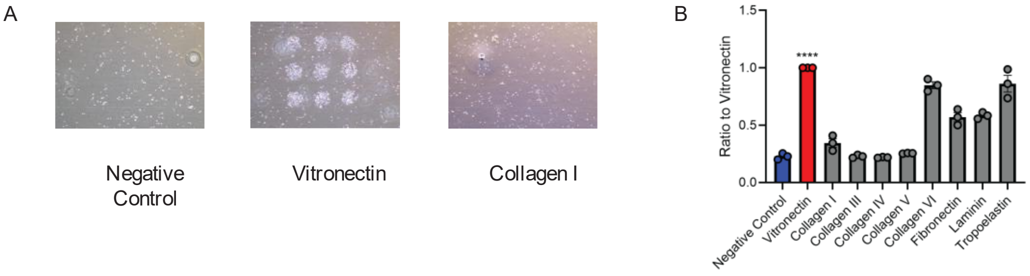


Bone marrow-derived macrophages (BMDMs) were cultured on an ECM Select Array. **(A)** Representative images of cell attachment with selective ECM conditions (negative control, vitronectin, and collagen I). **(B)** Quantification of cell attachment with different ECM conditions. ****, $p < 0.0001$. Values are shown as mean ± S.E.M. Two-tailed Welch's *t*-test was utilized.

**Supplementary Figure 2: Stiffness-activated macrophages have enhanced mitochondrial energetics.**

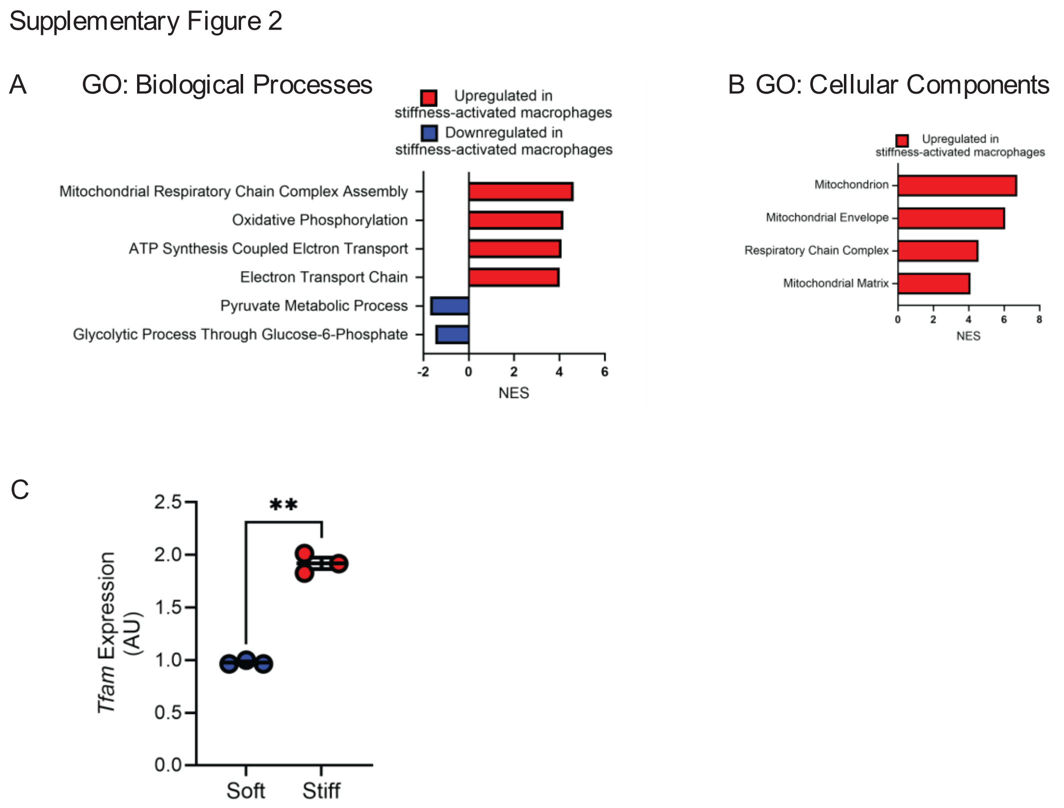


Bone marrow-derived macrophages (BMDMs) were cultured on soft or stiff matrix for 48 hr. mRNA was isolated and bulk RNA-seq was performed, and gene set enrichment analysis (GSEA) was performed using Gene Ontology (GO) Biological Processes and GO Cellular Components pathway compendia. Significantly regulated **(A)** GO Biological Processes and **(B)** GO Cellular Components were shown. **(C)** *Tfam* expression was measured by RT-PCR. **, $p < 0.01$. Values were shown as mean ± S.E.M.

**Supplementary Figure 3: Stiffness-activated macrophages have augmented glutaminolysis.**

Supplementary Figure 3

A

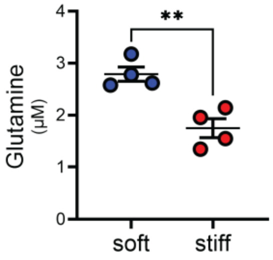


**(A)** Bone marrow-derived macrophages (BMDMs) were cultured on soft or stiff matrix for 48 hr. Intracellular glutamine levels were measured biochemically. **, $p < 0.01$. Values shown as mean ± S.E.M. Two-tailed Welch's $t$-test was utilized.

**Supplementary Figure 4: Enhanced glutaminolysis regulates macrophage apoptosis.**

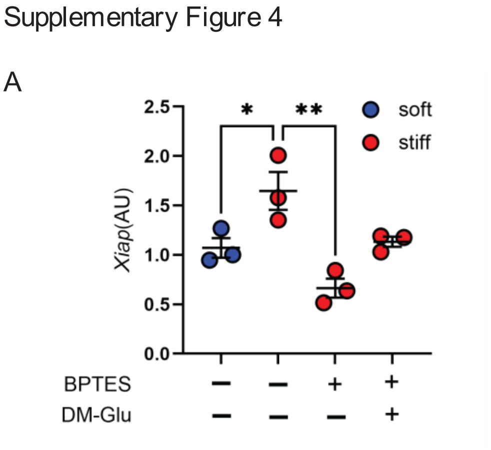


**(A)** Bone marrow-derived macrophages (BMDMs) were cultured on soft or stiff matrix in the presence or absence of the GLS inhibitor BPTES (3 μM) and DM-Glu (5 mM) for 48 hr. *Xiap* expression was measured by RT-PCR. *, $p < 0.05$; **, $p < 0.01$. Values were shown as mean ± S.E.M. One-way ANOVA followed by Tukey's multiple comparison test was utilized.
